# Redesigning methionyl-tRNA synthetase for *β*-methionine activity with adaptive landscape flattening and experiments

**DOI:** 10.1101/2022.12.28.522074

**Authors:** Vaitea Opuu, Giuliano Nigro, Christine Lazennec-Schurdevin, Yves Mechulam, Emmanuelle Schmitt, Thomas Simonson

**Affiliations:** Laboratoire de Biologie Structurale de la Cellule (CNRS UMR7654), Ecole Polytechnique, Institut Polytechnique de Paris, Palaiseau, France

**Author notes:** These authors contributed equally (Vaitea Opuu, Giuliano Nigro).

## Abstract

Amino acids (AAs) with a noncanonical backbone would be a valuable tool for protein engineering, enabling new structural motifs and building blocks. To incorporate them into an expanded genetic code, the first, key step is to obtain an appropriate aminoacyl-tRNA synthetase (aaRS). Currently, directed evolution is not available to optimize such AAs, since an appropriate selective pressure is not available. Computational protein design (CPD) is an alternative. We used a new CPD method to redesign MetRS and increase its activity towards *β*-Met, which has an extra backbone methylene. The new method considered a few active site positions for design and used a Monte Carlo exploration of the corresponding sequence space. During the exploration, a bias energy was adaptively learned, such that the free energy landscape of the apo enzyme was flattened. Enzyme variants could then be sampled, in the presence of the ligand and the bias energy, according to their *β*-Met binding affinities. Eleven predicted variants were chosen for experimental testing; all exhibited detectable activity for *β*-Met adenylation. Top predicted hits were characterized experimentally in detail. Dissociation constants, catalytic rates, and Michaelis constants for both *α*-Met and *β*-Met were measured. The best mutant retained a preference for *α*-Met over *β*-Met; however, the preference was reduced, compared to the wildtype, by a factor of 29. For this mutant, high resolution crystal structures were obtained in complex with both *α*-Met and *β*-Met, indicating that the predicted, active conformation of *β*-Met in the active site was retained.

**Author summary:** Amino acids (AAs) with a noncanonical backbone would be valuable for protein engineering, enabling new structural motifs. To incorporate them into an expanded genetic code, the key step is to obtain an appropriate aminoacyl-tRNA synthetase (aaRS). Currently, directed evolution is not available to optimize such AAs. Computational protein design is an alternative. We used a new method to redesign MetRS and increase its activity towards *β*-Met, which has an extra backbone methylene. The method considered a few active site positions for design and used a Monte Carlo exploration of sequence space, during which a bias energy was adaptively learned, such that the free energy landscape of the apo enzyme was flattened. Enzyme variants could then be sampled, in the presence of the ligand and the bias energy, according to their *β*-Met binding affinities. Eleven predicted variants were chosen for experimental testing; all exhibited detectable *β*-Met adenylation activity. Top hits were characterized experimentally in detail. The best mutant had its preference for *α*-Met over *β*-Met reduced by a factor of 29. Crystal structures indicated that the predicted, active conformation of *β*-Met in the active site was retained.

## Introduction

Each aminoacyl-tRNA synthetase (aaRS) attaches a specific amino acid to a tRNA that carries the corresponding anticodon, establishing the genetic code [1]. The attachment involves two steps: the amino acid (AA) reacts first with ATP to form aminoacyl adenylate. Next, the adenylate reacts with tRNA. Several aaRSs have been engineered experimentally to accept noncanonical amino acids (ncAAs) as preferred substrates [2–6]. This is the key step to make the ncAA part of an expanded code [3, 6, 7]. The ncAA can then be genetically encoded and incorporated into proteins by the cellular machinery. Thus, TyrRS was redesigned to be specific for several ncAAs [3, 6] and MetRS was redesigned to prefer azidonorleucine [8]. Several hundred ncAAs have been introduced into expanded codes, mostly using directed evolution to obtain the appropriate aaRSs. However, all these ncAAs had standard backbones.

In contrast, ncAAs with non standard backbones, such as D-AAs or *β*-AAs, would be of great interest in protein engineering, opening the possibility of new structural motifs and building blocks. For example, *β*-AAs have an extra backbone methylene that increases backbone flexibility, alters helical propensities [9, 10], increases the distance between *α* carbons, provides resistance to proteases, and can lead to modified side chain orientations in loop or sheet regions. To incorporate such ncAAs into an expanded code, the first step is to obtain an appropriate aaRS. However, directed evolution of aaRSs is still a major difficulty. The diversity one can explore and the selective pressure one can apply are limited, so the enzymes evolved so far have been weakly-active [11, 12]. In many situations, directed evolution is impossible, as no appropriate selective pressure can be applied. Thus, some archaeal tRNAs are not orthogonal to *E. coli*, precluding common selection methods. Some natural aaRSs have detectable activity for the ncAA of interest, precluding effective counterselection of the original AA activity by common methods. Thus, directed evolution has never been used to obtain aaRSs that were active towards nonstandard backbones.

Computer simulations that mimic directed evolution are another route, through computational protein design (CPD). Thus, tyrosyl-tRNA synthetase (TyrRS) was engineered recently to prefer the substrate D-Tyr over L-Tyr [13], using CPD to suggest mutations that were then tested experimentally. Recently, we engineered MetRS by CPD to obtain new variants with activity for adenylate formation by the native substrate *α*-Met, as a proof of principle [14]. Here, we turn to *β*-Met, an ncAA with a nonstandard backbone. We report the redesign of MetRS to decrease its preference for *α*-Met over *β*-Met, using a combination of CPD and experiments. The CPD calculations used a novel adaptive landscape flattening method [15, 16], which allows protein variants to be sampled according to their substrate binding affinity. A straightforward extension samples according to the binding free energy difference between two substrates [14, 16], such as *α*-Met and *β*-Met. This contrasts with most previous CPD work, where enzyme mutations were sampled according to the total energy of the enzyme-substrate complex [12, 13, 17–20].

To sample mutations that enhance ligand binding, we first flatten the free energy landscape of the apo enzyme in sequence space, using an adaptive, Wang-Landau Monte Carlo (MC) procedure to optimize a bias function that depends on sequence [15, 21]. The optimized bias approximates the free energy of the apo system, as a function of sequence, with its sign changed. Next, we simulate the protein–ligand complex, including the bias, which “subtracts out” the apo state. Remarkably, protein variants are then populated according to the apo/holo free energy difference, which is the binding free energy. Thus, the new method selects variants directly for substrate binding, and tight binders are exponentially enriched over the course of the MC trajectory. A straightforward extension allows us to sample for the *α*-Met/*β*-Met binding free energy difference, or specificity. The new methods were successful in the previous MetRS redesign [14], and we expected they would yield more hits and fewer false positives in the present application, compared to the older methods that considered the total energy [12, 13, 17–19].

Three positions close to the *β*-Met substrate were chosen for design. The CPD procedure sampled several thousand sequence variants. 18 predicted hits were tested, 10 of which exhibited detectable activity for *β*-MetAMP formation. The top four were characterized experimentally in considerable detail. The reactions of *α*-Met and *β*-Met with ATP to form the aminoacyl adenylates *α*-MetAMP and *β*-MetAMP were characterized by their catalytic efficiencies—the ratio between the rate constant for the catalytic step, *k*_cat_, and the Michaelis constant, *K*_*M*_. Dissociation constants, catalytic rates, and Michaelis constants for both *α*-Met and *β*-Met were measured. The best mutant retained a preference for *α*-Met over *β*-Met; however, the preference was reduced compared to the wildtype by a factor of 29. For this and one other mutant, high resolution crystal structures were obtained, both alone and in complex with *β*-Met. The increase in relative *β*-Met activity should considerably facilitate its further optimization using experimental directed evolution, once an effective selective pressure has been established. Indeed, directed evolution is more likely to succeed if the starting activity level (*β*-Met vs. *α*-Met) is not too low. More generally, the new method should facilitate the redesign of aaRSs and enzymes in general, and help expand the genetic code to include nonstandard backbone chemistries.

## Materials and methods

### Designing for ligand binding

The protein is modeled with molecular mechanics, with a fixed backbone and a discrete rotamer library for side chains, while the solvent is modeled implicitly (see below). We perform a MC exploration of the protein [16] with either no ligand (apo state), or a ligand, say L (holo state). We gradually increment a bias potential until all the side chain types at the mutating positions have roughly equal populations, thus flattening the free energy landscape. We number the mutating positions arbitrarily 1, …, *p*. The bias *E*^*B*^ at time *t* has the form:

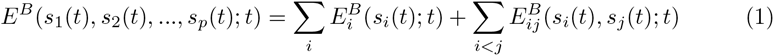

Here, *s*_*i*_(*t*) represents the side chain type at position *i*. The first sum is over single amino acid positions; the second is over pairs. The individual terms are updated at regular intervals of length *T*. At each update, whichever sequence variant (*s*_1_(*t*), *s*_2_(*t*), …, *s*_*p*_(*t*)) is populated is penalized by adding an increment 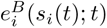 or 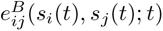 to each corresponding term in the bias. The increments have the form:

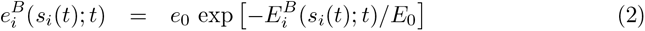

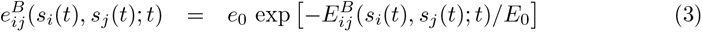

where *e*_0_ and *E*_0_ are constant energies. Over time, the bias for the most probable states grows until it pushes the system into other regions of sequence space.

The sampled population of a sequence *S* is normalized to give a probability, denoted 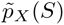, where *X* indicates which protein state is considered, apo or holo. The probability can be converted into a free energy 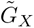 :

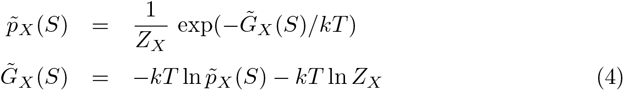

where *Z*_*X*_ is a normalization factor that depends on *X* but not *S*. We also have a relation between the free energies with and without the bias:

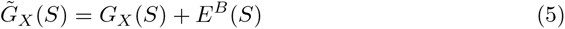

In practice, we will flatten the landscape not only of the apo state, but also the landscapes in the presence of the wildtype ligand and the new ligand targeted by the design. We will then take free energy differences between the three states, to obtain binding free energies and free energy differences between sampled sequences.

### Energy function and matrix

The energy was computed using an MMGBLK (molecular mechanics + Generalized Born + Surface Area or Lazaridis-Karplus):

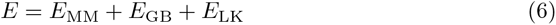

The MM term used the Amber ff99SB force field [22]. The GB term and its parametrization were described earlier [23]; similarly for the LK term [24]. The GB term used a “Fluctuating Dielectric Boundary” (FDB) method, where the GB interaction between two residues *I, J* was expressed as a polynomial function of their solvation radii [25]. These were kept up to date over the course of the MC simulation, so the GB interaction could be deduced on-the-fly with little additional calculation [25]. The solvent dielectric constant was 80; the protein dielectric was 6.8 [24].

To allow very fast MC simulations, we precomputed an energy matrix for each system [26, 27]. For each pair of residues *I, J* and all their allowed types and rotamers, we performed a short energy minimization (15 conjugate gradient steps) [16, 25]. The backbone was fixed (in its crystal geometry) and the energy only included interactions between the two side chains and with the backbone. At the end of the minimization, we computed the interaction energy between the two side chains. Side chain–backbone interaction energies were computed similarly (and formed the matrix diagonal).

### Structural models

#### MetRS complexes

Several MetRS complexes were modeled earlier and used here. For MetRS–ATP [14], we started from a crystal complex (PDB code 1PG0) between *E. coli* MetRS and a methionyl adenylate (MetAMP) analogue [28] with the KMSKS loop in its inactive conformation. Next, 15 loop residues from the the active loop conformation were transplanted from the *Leishmania major* structure (PDB code 3KFL) [29] by aligning common ligand fragments in both structures. Several loop residues were mutated into the *E. coli* types with Scwrl4 [30]. We adjusted the model geometry using 40 steps of conjugate gradient minimization to obtain a model of *E. coli* MetRS with the KMSKS loop in the active conformation. Finally, adenylate and pyrophosphate fragments were used to align ATP in the binding site. A Mg^2+^ ion was already in the 3KFL structure and was transferred to the new model. We call this model the MetRS–ATP complex. *α*-Met was added to form a MetRS–Met–ATP complex. Similarly, MetAMP was added, by superimposing it on the analogue present in 1PG0.

Recently [16], we added *β*-Met to MetRS–ATP. We used the recent crystal structure (PDB code 6SPN) of a complex between *E. coli* MetRS and *β*-Met [31]. We aligned that structure to MetRS–ATP to form MetRS–*β*-Met–ATP. The 6SPN crystal structure included two conformations for the *β*-Met carboxylate. One is much closer to the ATP, as required for catalysis, and we chose that geometry. Complexes with the activated substrates [*α*-MetATP]^‡^] and [*β*-Met–ATP]^‡^] were built earlier [14, 16], using a restrained minimization to produce the appropriate pentavalent geometry around the ATP *α* phosphorus. Here, we also formed the complex with *β*-MetAMP, by superimposing the latter onto MetAMP in the MetRS–MetAMP complex. The complex with *β*-ValAMP was constructed by replacing the amino acid moiety.

For each complex, the geometry of the protein around the ligands was relaxed slightly by performing a short, restrained molecular dynamics simulation, with the ligands held fixed. The entire system was placed in a large box of explicit TIP3P water [32]. Harmonic restraints were applied to nonhydrogen atoms, with force constants that decreased gradually from 5 to 0.5 kcal/mol/Å^2^ over 575 ps of dynamics, performed with the NAMD program [33]. The final protein geometry was used for the design calculations. We have found that this procedure reduces specialization of the backbone model towards each particular ligand.

For the design calculations, the protein backbone and side chains more than 20 Å from the ligand were held fixed. The other side chains were allowed to explore rotamers, taken from the Tuffery library, augmented to allow multiple orientations for certain hydrogen atoms [16, 34]. For *α*-Met and *β*-Met, we allowed the Met rotamers from the Tuffery library, with the rest of the ligand held fixed. Side chains 13, 256 and 297 were allowed to mutate into all types except Gly and Pro. Thus, there were 5832 possible sequences in all. Histidine protonation states at non-mutating positions were assigned by visual inspection of the 3D structure. System preparation was done using the protX module of the Proteus design software [35].

#### Unfolded state

The unfolded state energy was estimated with a tri-peptide model [36]. For each mutating position, side chain type, and rotamer, we computed the interaction between the side chain and the tri-peptide it forms with the two adjacent backbone and C_*β*_ groups. Then, for each allowed type, we computed the energy of the best rotamer and averaged over mutating positions. The mean energy for each type was taken to be its contribution to the unfolded state energy. The contributions of the mutating positions were summed to give the total unfolded energy.

### Ligand force field

The partial charges of ribose, adenine and side chain fragments were derived from existing Amber parameters in analogous fragments. For the junction atoms between *β*-Met and AMP, we performed an HF/6-31G* ab initio calculation with Gaussian 9, and partial charges were chosen to reproduce the electrostatic potential, following the usual Amber procedure [22]. Bonded and van der Waals parameters were assigned by analogy to the *α*-Met model [14]. Parameters for the implicit solvent energy terms were assigned by analogy to existing groups. The Mg charge was set to +1.5, as previously [14].

### Monte Carlo simulations

To optimize the bias potential, we performed MC simulations of the considered state with bias updates every *T* = 1000 steps, with *e*_0_ = 0.2 kcal/mol and *E*_0_ = 50 kcal/mol [15]. During the first 10^8^ MC steps, we optimized a bias potential including only single-position terms. There were *p* = 3 mutating positions, which all contributed to the bias. In the second stage, we ran MC simulations of 5.10^8^ MC steps [16], using 8 replicas with thermal energies (kcal/mol) of 0.17, 0.26, 0.39, 0.59, 0.88, 1.33, 2.0 and 3.0. Temperature swaps were attempted every 500 steps. All the replicas experienced the same bias potential. Both stages used 1- and 2-position moves.

### Experimental mutagenesis and kinetic assays

#### Purification of wildtype and mutant MetRS

Throughout this study, we used a His-tagged M547 monomeric version of *E. coli* MetRS, fully active, both *in vitro* and *in vivo* [37]. The gene encoding M547 MetRS from pBSM547+ [38, 39] was subcloned into pET15blpa [40] to overproduce the His-tagged enzyme in *E. coli* [31]. Site-directed mutations were generated using the QuikChange method [41], and the whole mutated genes verified by DNA sequencing. The enzyme and its variants were produced in BLR(DE3) *E. coli* cells. Transformed cells were grown overnight at 37°C in 0.25 L of TBAI autoinducible medium containing 50 *μ*g/ml ampicillin. They were harvested by centrifugation and resuspended in 20 ml of buffer A (10 mM Hepes-HCl pH 7.0, 3 mM 2-mercaptoethanol, 500 mM NaCl). They were disrupted by sonication (5 min, 0°C), and debris was removed by centrifugation (15,300 G, 15 min). The supernatant was applied on a Talon affinity column (10 ml; Clontech) equilibrated in buffer A. The column was washed with buffer A plus 10 mM imidazole and eluted with 125 mM imidazole in buffer A. Fractions containing tagged MetRS were pooled and diluted ten-fold in 10 mM Hepes-HCl pH 7.0, 10 mM 2-mercaptoethanol (buffer B). These solutions were applied on a Q Hiload ion exchange column (16 mL, GE-Healthcare), equilibrated in buffer B containing 50 mM NaCl. The column was washed with buffer B and eluted with a linear gradient from 5 to 500 mM NaCl in buffer B (2 ml/min, 10 mM/min). Fractions containing tagged MetRS were pooled, dialyzed against a 10 mM Hepes-HCl buffer (pH 7.0) containing 55% glycerol, and stored at -20°C. The MetRS was estimated by SDS-PAGE to be at least 95% pure.

#### Measurement of ATP-PPi isotopic exchange activity

Prior to activity measurements, MetRS was diluted in standard buffer (20 mM Tris-HCl buffer pH 7.6, 10 mM 2-mercaptoethanol, 0.1 mM EDTA) containing 0.2 mg/ml bovine serum albumin (Aldrich) if the concentration after dilution was less than 1 *μ*M. Initial rates of ATP-PPi exchange activity were measured at 25°C as described [42]. In brief, the 100 *μ*l reaction mixture contained Tris-HCl (20 mM, pH 7.6), MgCl2 (7 mM), ATP (2 mM), [^32^P]PPi (1800-3700 Bq, 2 mM) and various concentrations (0-16 mM) of the Met amino acid. The exchange reaction (Fig 1) was started by adding catalytic amounts of MetRS (20 *μ*l). After quenching the reaction, ^32^P-labeled ATP was adsorbed on charcoal, filtered, and measured by scintillation counting. For *β*-Val-dependent activity measurements, *α*-Met was replaced by 10 mM *β*-Val. In this case, all reaction components were treated with MGL to remove contaminating *α*-Met.

**Fig 1.**
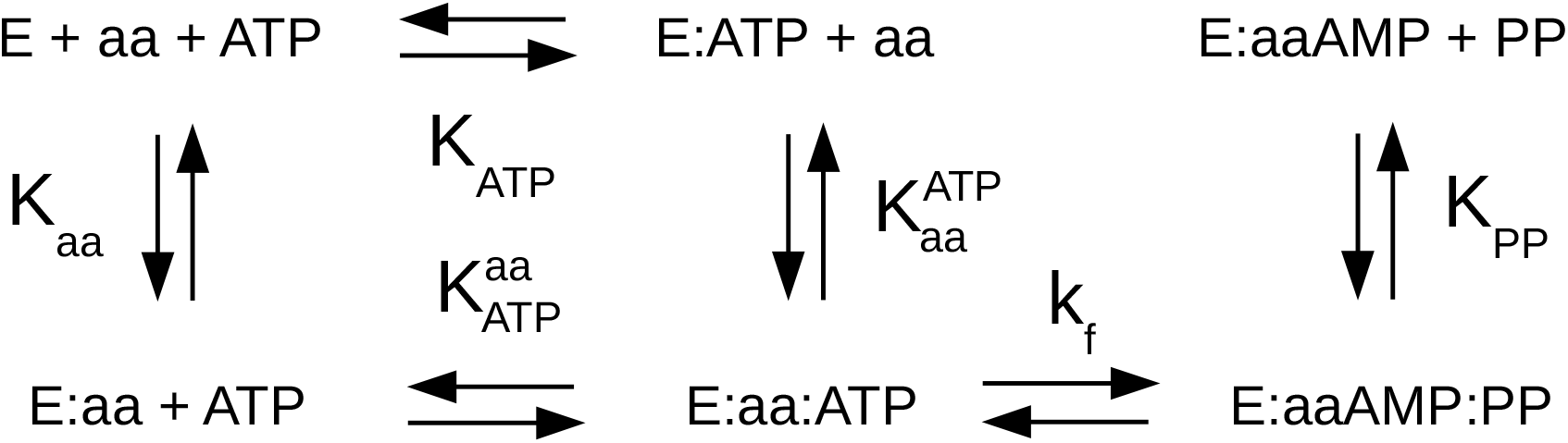
MetRS reactions to form aminoacyl adenylate. The amino acid is denoted aa and the enzyme E. Equilibrium constants are in uppercase; *k*_*f*_ is a rate constant.

#### Measurement of ATP-PPi exchange activity

Prior to activity measurements, MetRS or its variants were diluted in standard buffer (20 mM Tris-HCl buffer pH 7.6, 10 mM 2-mercaptoethanol, 0.1 mM EDTA) containing 0.2 mg/ml bovine serum albumin (Aldrich) if the concentration after dilution was less than 1 *μ*M. Initial rates of ATP-PPi exchange activity were measured at 25°C as described [31]. In brief, the 100 *μ*l reaction mixture contained Tris-HCl (20 mM, pH 7.6), MgCl2 (7 mM), ATP (2 mM), [32P]PPi (1800-3700 Bq, 2 mM) and various concentrations of *α*-Met. The exchange reaction (Fig 1) was started by adding catalytic amounts of MetRS (20 *μ*l). After quenching the reaction, 32P-labeled ATP was adsorbed on charcoal, filtered, and measured by scintillation counting. For *β*-Val-dependent activity measurements, *α*-Met was replaced by 10mM *β*-Val. In this case, all reaction components were treated with MGL to remove contaminating *α*-Met.

#### Fluorescence at equilibrium

Variations of the intrinsic fluorescence of M547 and its variants (0.5 *μ*M) upon titration with substrates were followed at 25°C in 20 mM Tris-HCl (pH7.6), 10 mM 2-mercaptoethanol, 2 mM MgCl2 and 0.1mM EDTA as described [31, 42, 43]. Measurements were done in a Hellma 1 cm *×* 0.4 cm cuvette with an FP-8300 JASCO spectrofluorometer (295 nm excitation, 340 nm emission). All titration curves were corrected for dilution. Concentrations of *α*-Met or *β*-Met were varied from 3 *μ*M to 1 mM and from 0.06 mM to 4 mM, respectively. Data were fitted to simple saturation curves from which the corresponding dissociation constants were derived using the Origin software (OriginLab Corp.).

#### Fluorescence at the pre-steady state

Fluorescence measurements at the pre-steady-state were performed as described [31, 42, 44] using an SX20 stopped flow apparatus (Applied Photophysics, UK). All experiments were performed in standard buffer supplemented with 2 mM MgCl2. The formation of *α*- or *β*-methionyl adenylate was initiated by mixing 1:1 (v/v) an enzyme solution (1 *μ*M for *α*-Met or 2 *μ*M for *β*-Met) containing ATP-Mg^2+^ (2 mM) and PPi (10 *μ*M) with a solution containing the same concentrations of ATP-Mg^2+^ plus variable amounts of the amino acid (10 *μ*M to 640 *μ*M for *α*-Met or 25 *μ*M to 2 mM for *β*-Met). After mixing, fluorescence was recorded and fitted to single exponentials from which the rate constants were derived. Each rate was determined three times. Kinetic (k_*f*_ (aa)) and equilibrium 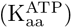 parameters are defined in Fig 1. They were deduced from the fit of the measured rate constants to the theoretical saturation curves [42–44] using Origin. Each experiment was performed at least twice independently. Results below are expressed as mean *±* either standard deviation from the independent experiments or standard error from the fitting procedure, whichever is greater. All experiments were performed using MGL-treated *β*-Met.

### X-ray structure determination

Crystals of the MetRS CAC and MAC variants were obtained by microseeding with crystals of M547 [45] in a solution containing 30 mM KPO_4_, 4 mM 2-mercaptoethanol, 1.08 mM ammonium citrate (pH 7.0) and 3.6 mg/mL of protein. Type 2 crystals [37] of the apo-CAC and -MAC enzymes were chosen. For the structures of MetRS:*β*-Met complexes, 10 mM *β*-Met and 1 mM adenosine were added to the crystallization medium prior to microseeding. Before data collection, crystals were quickly soaked in a solution containing 1.4 M ammonium citrate, 30 mM potassium phosphate (pH 7.0) and 25% v/v of glycerol and flash-cooled in liquid nitrogen. In the case of MetRS:*β*-Met complexes, the cryoprotecting solution was supplemented with 10 mM *β*-Met. Data were collected at the Proxima 2 beamline at the SOLEIL synchrotron (Gif sur Yvette, France). Diffraction images were analyzed with the XDS program [46] and the data further processed using programs from the CCP4 package [47]. The structure was solved by rigid body refinement of the wild-type MetRS model (PDB id 3H9C) [37], using PHENIX [48]. Coordinates and associated B factors were refined through several cycles of manual adjustments with Coot [49] and positional refinement with PHENIX. For MetRS:*β*-Met structures, no bound adenosine was observed. Data collection and refinement statistics are summarized in Supplementary Table SM1. Attempts to obtain the structures of the two variants bound to *α*-Met only revealed a very low occupancy of the amino acid binding site and were not analyzed further.

## Results

### Computational design of MetRS to bind *β*-MetAMP and *β*-ValAMP

MetRS was redesigned through adaptive MC simulations where three active site positions, 13, 256, 297 (Fig 2) were allowed to mutate freely. With *β*-MetAMP binding as the design target, adaptive landscape flattening was applied to the apo state. In practice, over the course of a long MC simulation, the bias was gradually optimized, until the free energy landscape was sufficiently flattened (all amino acid types were sampled at all three positions with roughly equal populations) [15]. Then, a second simulation was done, of the holo state, with the bias included. Sequences sampled with either *α*-MetAMP or *β*-MetAMP as the ligand are shown in Fig 3, as logos. With the *α*-Met ligand, the wildtype amino acid types, L, A, I were highly ranked at all three designed positions.

**Fig 2.**
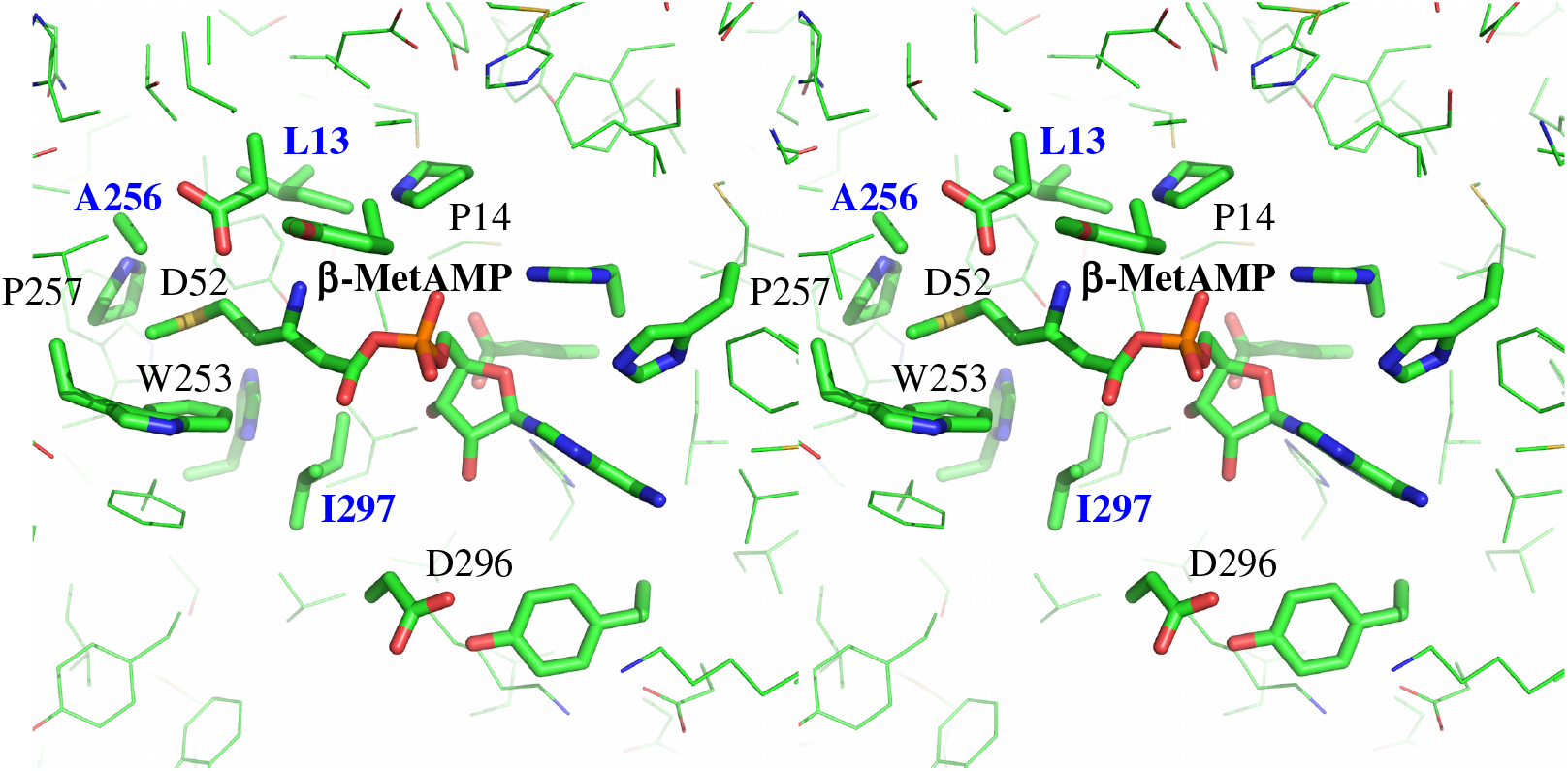
Stereo view of the wildtype MetRS binding pocket, showing *β*-MetAMP and selected side chains.

**Fig 3.**
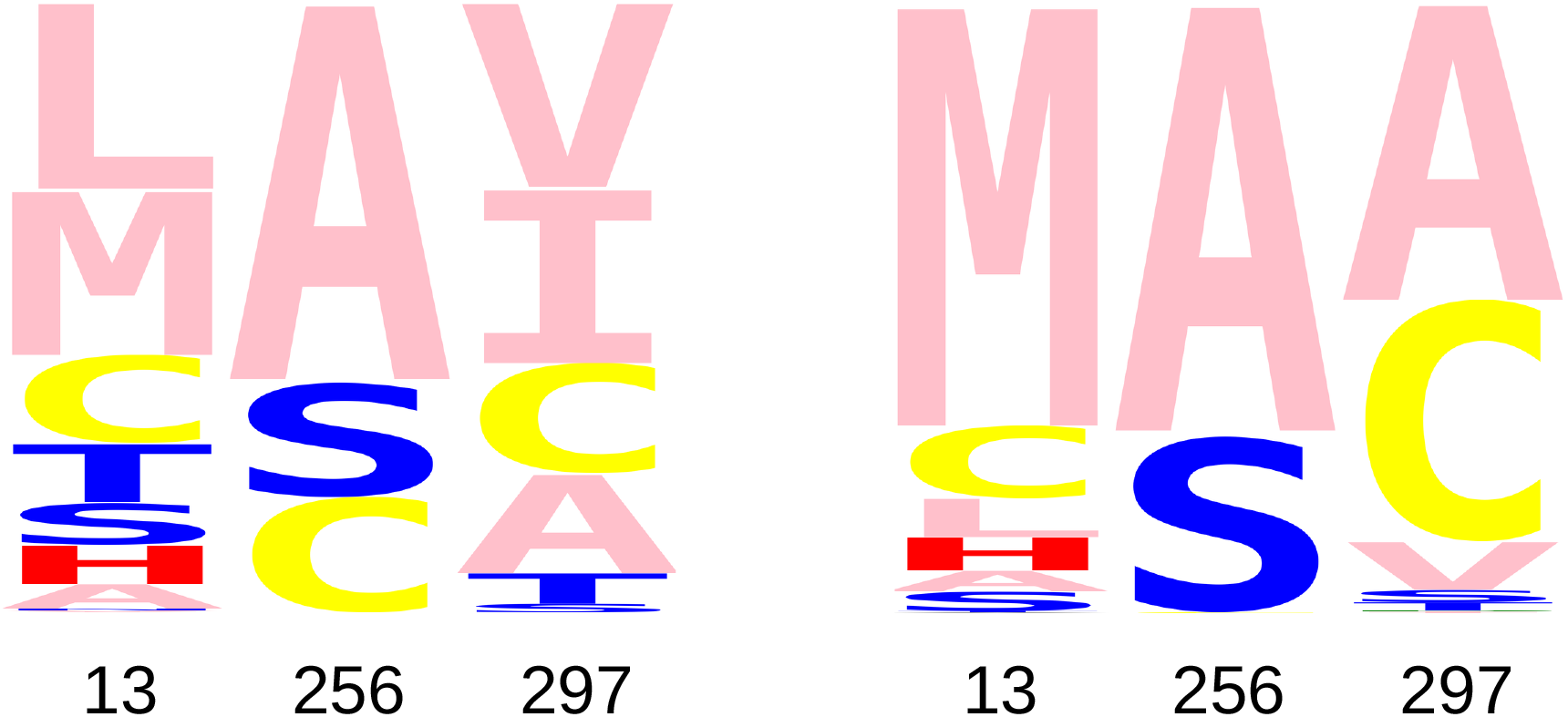
Designed sequence logos. Left) MetRS sequences sampled according to their predicted *α*-MetAMP binding free energies. The three mutating positions, 13, 256, 297 are shown. The height of each letter measures the frequency of its type, when the sampled sequences are weighted by their ligand binding free energies. **Right)** MetRS sequences sampled according to their predicted *β*-MetAMP binding free energies.

With *β*-Met binding as the design criterion, the wildtype variant LAI was not sampled in the holo state. However, the similar variants LSI, MAI and LAA were sampled. Their predicted binding free energies were within 0.5 kcal/mol of each other and can be taken as points of reference. 35 variants were predicted to have *β*-MetAMP binding that was improved, compared to LSI. 25 were improved by 0.5 kcal/mol or more. The maximum improvement was 3.6 kcal/mol. The predominant amino acid types (Fig 3, right) were MCL at position 13, AS at position 256, and ACV at position 297. The top 10 predictions are listed in Table 1. Several predicted variants were active, as detailed in the next section.

**Table 1.**
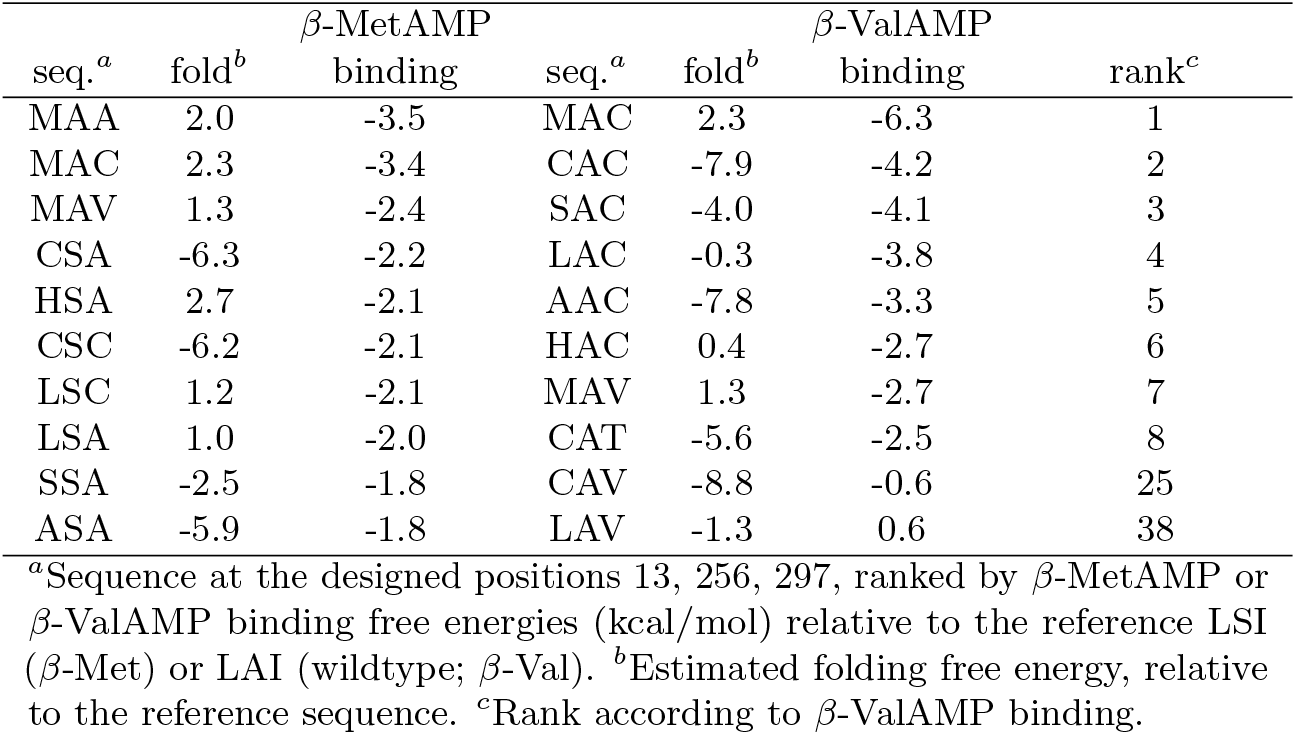
Predicted *β*-MetAMP and *β*-ValAMP binding affinities for top MetRS redesigns

| β-MetAMP | | | β-ValAMP | | | |
| --- | --- | --- | --- | --- | --- | --- |
| seq.<sup>a</sup> | fold<sup>b</sup> | binding | seq.<sup>a</sup> | fold<sup>b</sup> | binding | rank<sup>c</sup> |
| MAA | 2.0 | -3.5 | MAC | 2.3 | -6.3 | 1 |
| MAC | 2.3 | -3.4 | CAC | -7.9 | -4.2 | 2 |
| MAV | 1.3 | -2.4 | SAC | -4.0 | -4.1 | 3 |
| CSA | -6.3 | -2.2 | LAC | -0.3 | -3.8 | 4 |
| HSA | 2.7 | -2.1 | AAC | -7.8 | -3.3 | 5 |
| CSC | -6.2 | -2.1 | HAC | 0.4 | -2.7 | 6 |
| LSC | 1.2 | -2.1 | MAV | 1.3 | -2.7 | 7 |
| LSA | 1.0 | -2.0 | CAT | -5.6 | -2.5 | 8 |
| SSA | -2.5 | -1.8 | CAV | -8.8 | -0.6 | 25 |
| ASA | -5.9 | -1.8 | LAV | -1.3 | 0.6 | 38 |

<sup>a</sup>Sequence at the designed positions 13, 256, 297, ranked by $\beta$ -MetAMP or $\beta$ -ValAMP binding free energies (kcal/mol) relative to the reference LSI ( $\beta$ -Met) or LAI (wildtype; $\beta$ -Val). <sup>b</sup>Estimated folding free energy, relative to the reference sequence. <sup>c</sup>Rank according to $\beta$ -ValAMP binding.

Redesign for *β*-Val activity was unsuccessful. We targeted *β*-ValAMP binding, varying the same three positions. The top 8 predictions are included in Table 1. In the top variants, the native A was predominant at position 256, while position 297 strongly preferred C and position 13 was more variable. Experimental tests were done for four mutants: CAC and LAC, among the top predictions, and CAV and LAV, which were ranked somewhat lower. No *β*-Val-dependent ATP-PPi isotopic exchange activity was detected.

### Experimental characterization of four variants designed for *β*-Met activity

From the 35 top predicted variants, 18 were chosen for experimental testing, such that they recapitulated the predominant amino acid types predicted at each redesigned position. They were tested for *β*-MetAMP synthesis using the pre-steady state fluorescence assay in the presence of 2 mM ATP-Mg^2+^ and 10 mM *β*-Met. All reaction components were extensively treated with MGL, to remove any contaminating *α*-Met. 10 of the 18 variants were active, albeit with a reduced, not an increased activity compared to the wildtype. Variants that included an A256S mutation were all inactive. The others all preserved the native Ala256, associated with A, C, V, or native I at position 297 and M, C, L or S at position 13. Details and experimental kinetic parameters are reported in Supplementary Table SM2. Below, we report a detailed characterization of the four best experimental variants: MAC, CAC, SAC, and LAC, which are all compared to the wildtype LAI. “Best” variants were chosen for their relative *α*-Met and *β*-Met activities, reported in Table SM2.

### Selected mutations at positions 13 and 297 strongly affect MetAMP formation

The enzyme reaction constants are defined in Fig 1. Experimental values are reported in Table 2. The *α*-Met dissociation constants K_*d*_ of the mutants were increased 3-4 fold, based on steady-state fluorescence measurements. With ATP present, the *α*-Met dissociation constants 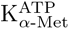 were increased more strongly, by factors of 15 and 19 for MAC and LAC, and by factors of 30 for CAC and SAC, based on the fluorescence at the pre-steady state. We also derived the rate constants *k*_*f*_ for *α*-Met activation. The rates were decreased by an order of magnitude for MAC and by a factor of 2 for LAC.

**Table 2.** Experimental parameters for top MetRS redesigns

|  | LAI (WT) | MAC | CAC | SAC | LAC |
| --- | --- | --- | --- | --- | --- |
| $K_d(\alpha\text{-Met})$ (mM) | 0.050(5)<sup>c</sup> | 0.16(2) | 0.17(3) | 0.17(2) | 0.19(3) |
| $K_{\alpha\text{-Met}}^{\text{ATP}}$ (mM) | 0.070(4) | 1.1(2) | >5 | >5 | 1.3(1) |
| $k_f(\alpha\text{-Met})$ (s<sup>-1</sup>) | 260±10 | 33.9±1.9 | NA | NA | 139±17 |
| $k_f(\alpha\text{-Met})/K_{\alpha\text{-Met}}^{\text{ATP}}$ (s<sup>-1</sup>mM<sup>-1</sup>) | 3714±143 | 31.5±4.4 | 14.7±0.1 | 17.9±1.5 | 107±2 |
| aminoacylation rate<sup>a</sup> (s<sup>-1</sup>) | 1.9(2) | 0.017(4) | 0.013(2) | 0.016(3) | 0.07(1) |
| $K_d(\beta\text{-Met})$ (mM) | 1.2(2) | 0.9(3) | 0.9(2) | 1.0(3) | 0.9(3) |
| $K_{\beta\text{-Met}}^{\text{ATP}}$ (mM) | 0.26 | 0.58 | 0.56 | 1.3 | 0.26 |
| $k_f(\beta\text{-Met})$ (s<sup>-1</sup>) | 0.047(4) | 0.0083(3) | 0.0105(8) | 0.007(1) | 0.013(1) |
| $k_f(\beta\text{-Met})/K_{\beta\text{-Met}}^{\text{ATP}}$ (s<sup>-1</sup>mM<sup>-1</sup>) | 0.180(4) | 0.015(3) | 0.021(8) | 0.0054(12) | 0.05(1) |
| $10^4 \times \beta\text{-Met}/\alpha\text{-Met}$ ratio<sup>b</sup> | 0.5 | 5.0 | 14.0 | 3.0 | 5.0 |
| β-Met enhancement | 1 | 10 | 29 | 6 | 10 |

Experimental enzyme parameters. <sup>a</sup>Initial rate for tRNA acylation with $\alpha$ -Met, measured as reported earlier [31]. <sup>b</sup> $k_f(\text{aa})/K_{\text{aa}}^{\text{ATP}}$ ratio. <sup>c</sup>In parentheses: experimental uncertainty in significant digits.

For CAC and SAC, the rates could not be measured, as 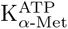 was too large. Catalytic efficiencies 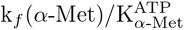 for *α*-Met activation were reduced by two orders of magnitude for MAC, CAC and SAC, and by a factor of more than 30 for LAC. Note that this parameter directly reflects the free energy difference between the transition state complex and the E:ATP:Mg^2+^ + *α*-Met state. These reduced efficiencies for the activation reaction are also reflected in the off rates of the global aminoacylation reaction, which were reduced in similar proportions. Overall, the *α*-Met catalytic efficiencies were reduced (transition state binding was weakened) and the mutations at both positions (13 and 297) contributed (the single mutant LAC was more active than the double mutants). Reduction was largest for the CAC and SAC variants (Table 2).

### Activation of *β*-Met is much less affected than that of *α*-Met

The *β*-Met used throughout was treated enzymatically to remove any contaminating *α*-Met. The enzyme employed was a bacterial methionine *γ*-lyase, which breaks down *α*-Met but not *β*-Met [31]. From steady-state fluorescence measurements, the *β*-Met dissociation constants of the mutants were only slightly reduced, compared to wildtype MetRS (25% decrease; Table 2). On the other hand, with ATP present, pre-steady-state fluorescence showed that the dissociation constants 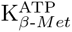 were unaffected in LAC, increased by a factor of 2 in MAC and SAC, and by a factor of 5 in SAC. The rates k_*f*_ (*β*-Met) of *β*-Met activation were decreased by factors ranging from 4 (LAC) to 7 (SAC). The catalytic efficiency was reduced by a factor of 3.6 for the single mutant (LAC). For the double mutants, the lowering of the catalytic efficiency was greatest for CAC (8.6 fold). In all cases, activation of *β*-Met was less affected than that of *α*-Met. As a result, the selectivity of all four mutant enzymes for *β*-Met vs. *α*-Met was increased. For SAC, LAC and MAC, the increase was between 6 and 10. For the double mutant CAC, the *β*-Met vs. *α*-Met selectivity was increased 29-fold.

### Crystal structures of the CAC and MAC variants

The structures of the CAC and MAC variants were solved in both the apo and *β*-Met bound forms. The apo structures did not show any significant conformational changes compared to the wild-type enzyme (Supplementary Figure SM1 A-B). The substituted residues, C13 and C297 (CAC), or M13 and C297 (MAC) were clearly visible in the electron density. Additional electron density close to the sulfur atom of C297 was visible in both structures. We modelled this density as a water molecule, which refined at a distance from the sulfur atom of 2.6 Å (CAC) or 2.3 Å (MAC; Supplementary Figure SM1 C-D). Although the density was best accounted for by a Cys and a water molecule, it cannot be excluded that a fraction of C297 residues in the crystal have been oxidized to sulfenic acid. Indeed, X-ray induced oxidation of Cys has already been observed and proposed to require a reactive cysteine near a water molecule [50].

For both holo structures, *β*-Met was clearly visible in the active site. In wild-type MetRS, binding of *β*-Met and *α*-Met, both occur via a ligand-induced, concerted rearrangement of aromatic side chains (W229, W253, F300 and F304) [31] that puts W253 in contact with the Met side chain. This rearrangement did not fully occur with the present MetRS variants upon *β*-Met binding. With CAC, F300 and F304 remained in their apo positions whereas a minor fraction of W229 was rotated (Supplementary Figure SM2). W253 mainly rotated to a position that differs from that in wild-type MetRS:*β*-Met. With MAC, the rearrangement was closer to that observed in the wild-type enzyme. W253 and F304 mainly rotated to the same position as in the wild-type MetRS:*β*-Met complex, but only minor fractions of W229 and F300 moved away from their apo positions. These partial rearrangements in the CAC and MAC variants probably contribute to the observed loss of catalytic efficiency towards both *α*-Met and *β*-Met (Table 2).

In the wild-type MetRS:*α*-Met complex, the concerted rearrangement of aromatic residues is accompanied by a rotation of Y15 side chain that places the *α*-Met carboxylate in its active position (position 1), favorable to catalysis [28, 37, 51]. In the *β*-Met complex, the rotated position of Y15 was unstable and several alternative conformations were observed [31]. Concomitantly, two alternative positions of the *β*-Met carboxylate were observed. In the major conformation (position 2), the carboxylate pointed away from the active site, preventing the full rotation of Y15. In the minor conformation (position 3), the carboxylate was closer to the active state seen with *α*-Met (position 1).

With the CAC variant, the *β*-Met carboxylate mainly adopted a position (position 4) closer to position 1 than position 3 (Supplementary Fig SM1 E). Consistent with this, the locking conformation of Y15 was more highly occupied (Supplementary Fig SM2 A-B and D). Thus, in the CAC variant, the *β*-Met carboxylate is mainly in a position favorable to catalysis. This might account for the increased selectivity of this variant towards *β*-Met. With the MAC variant, in contrast, only position 2 of the *β*-Met carboxylate was visible (Supplementary Fig SM2 C).

The CAC and MAC variants both had a carbon at position 297 of the amino acid binding pocket. As mentioned above, C297 tightly bound a water molecule that was 4.5 Å away from the *β*-Met sulphur atom. This likely contributes to *β*-Met/*α*-Met binding. Interestingly, in the CAC variant, the C297 sulphur atom was 4.5 Å away from the *β*-Met carboxylate (Supplementary Fig SM2 A). Thus, C297 may favor the activation of *β*-Met more than that of *α*-Met.

In the CAC variant and the wild-type enzyme, the main chain oxygen of residue 13 interacted with the amino group of *β*-Met and the Y15 main chain nitrogen interacted with a *β*-Met carboxylate oxygen (Supplementary Fig SM2 B,C). Because of the bulkier Met side chain at position 13, the MAC variant showed a slight displacement of the main chain (0.5 Å for the C_*α*_ of residue 13). This displacement probably leads to subtle changes in the *β*-Met environment during catalysis. Overall, it appears that better positioning of the *β*-Met carboxylate and tuning of the conformation of the A12-Y15 region are good ways to enhance the selectivity of MetRS for *β*-Met over *α*-Met.

## Concluding discussion

Genetic code expansion for noncanonical backbones would open exciting new directions for protein engineering, allowing new structural motifs and building blocks. We envisage that ncAAs could be developed in two “orthogonal” directions, where a variety of noncanonical backbones could be combined with different noncanonical side chains, leading to a combinatorial space of ncAAs. Unfortunately, directed evolution of aaRSs is still a major difficulty, and has never been used to obtain activity towards noncanonical backbones. One difficulty is that directed evolution requires not only a selective pressure, but also a starting enzyme that has a certain level of activity towards the ncAA. Natural MetRS enzymes do not provide this.

Here, we used a mixed computational and experimental approach. We searched for *β*-Met activity using CPD, then tested the predictions experimentally. CPD exploration used a powerful new adaptive sampling method. Indeed, most previous redesign studies of protein-ligand binding used the total system energy as the design target [20]. While many successes have been reported, the predicted designs often had a low activity, and many false positives were produced [11–13]. Here, instead of targeting the total energy, we used the new, adaptive sampling [14, 15] to target ligand binding directly, applying positive design to the bound state and negative design to the unbound. 18 variants were selected for testing: 10 displayed detectable activity for *β*-Met.

The top four variants were characterized in detail experimentally. They had experimental preferences for *α*-Met that were reduced, relative to *β*-Met, by factors of 6, 10, 10, and 29. These reductions were due to reduced *α*-Met activity, whereas the *β*-Met activites were close to the wildtype level. The high resolution X-ray structure of the best mutants, CAC and MAC, showed that the AA position was close to that used in the CPD calculations, and the active carboxylate moiety was positioned as expected to react with ATP. While the CPD calculations only considered the binding of *β*-Met adenylate, the top variants were also shown to aminoacylate cognate tRNA with *β*-Met.

Although the CPD calculations overestimated the strength of the redesigned *β*-Met binding, they did not produce any false positives for the three positions desiged here. Additional calculations that targeted other positions did produce false positives, possibly because they considered positions that could affect the conformation of the flexible KMSKS loop in the active site [42]. Design for *β*-Val also produced false positives. This could be due to the use of an incorrect *β*-Val pose in the calculations; a more advanced study involving molecular dynamics (MD) and an explicit solvent model could be used to test this possibility. More sophisticated and expensive CPD procedures involving MD exploration of backbone degrees of freedom are another possibility [52, 53].

The new methodology used here is applicable to many problems. For MetRS, it was used earlier to retrieve variants with azidonorleucine (Anl) activity [14]. The top experimental variants were retrieved when the enzyme was redesigned for ligand binding *specificity* (Anl vs. Met), rather than Anl affinity. Specificity of transition state binding can also be targeted, in principle. The methodology was used here with a model where backbone flexibility was treated implicitly through a dielectric continuum model, implemented within the Proteus software [35]. However, the methodology is general and could be combined with other methods and software, such as lambda-dynamics with the Charmm software [52]. The successful design of MetRS variants with a large decrease in *α*-Met activity, relative to *β*-Met, indicates that the assumptions used here are not too limiting, and that substrate binding affinity, as a design target, is a good proxy for activity. Experimental directed evolution is a future perspective, assuming an appropriate selective pressure for an extra backbone methylene can be identified. The best present design, with its 29-fold gain in relative *β*-Met activity, should be a valid starting point.

## Supporting information

**S1 Appendix. Additional data** are provided as a Supplementary Appendix, which provides statistics on X-ray data collection, experimental kinetic parameters for selected MetRS mutants, and 3D active site structure views for selected systems.

**S1 File. X-ray structures** are provided in four separate CIF files for the CAC and MAC mutants, determined with and without bound *β*-Met. The PDB identifiers are 8BRU (apo-MAC), 8BRV (MAC-*β*-Met), 8BRW (apo-CAC), 8BRX (CAC-*β*-Met).

